# Overall biomass yield on multiple nutrient sources

**DOI:** 10.1101/2023.02.16.528813

**Authors:** Ohad Golan, Olivia Gampp, Lina Eckert, Uwe Sauer

**Affiliations:** Institute of Molecular Systems Biology, ETH Zürich, Otto-Stern-Weg 3, 8093 Zürich, Switzerland; Life Science Zurich PhD Program on Systems Biology, Zurich, Switzerland

## Abstract

Microorganisms utilize nutrients primarily to generate biomass and replicate. When a single nutrient source is available, the produced biomass increases linearly with the initial amount of the available nutrient. This linear trend can be predicted to high accuracy by “black box models” that consider growth as a single chemical reaction with nutrients as substrates and biomass as a product. Since natural environments typically feature multiple nutrients, we here quantify the effect of co-utilization of multiple nutrients on bacterial biomass production. First, we demonstrate a mutual effect between the metabolism of different nutrient sources where the ability to utilize one is affected by the other. Second, we show that for some nutrient combinations, the produced biomass is no longer linear to the initial amount of nutrients. These observations cannot be explained by the traditional “black box models”, presumably because the metabolism of one nutrient affects another, which is not accounted for by these models. To capture these observations, we extent “black box models” to include catabolism, anabolism, and biosynthesis of biomass precursors and phenomenologically add a mutual effect between the metabolism of the nutrient sources. The expanded model qualitatively recaptures the experimental observations and, unexpectedly, predicts that the produced biomass is not only dependent on the combination of nutrient sources but also on their relative initial amounts. We validate this prediction experimentally by demonstrating how measurement of the produced biomass can be used to determine how each nutrient effects the metabolic processes of another.

## Introduction

Natural environments are characterized by a broad spectrum of physicochemical parameters that collectively define constraints within which species survive and thrive. Of particular importance to niche occupancy by different microbes are the type of nutrients and their temporal availability. For example, bacteria growing in a riverbed might experience continuous nutrient flux and high spatial homogeneity while bacteria growing in pulsating environments, such as tidal wetlands or at the sea bottom, receive nutrients only sporadically [1, 2]. Different physiological traits give a fitness advantage for different nutrient dynamics. In conditions of continuous nutrient flux, organisms with higher growth rate are favored while environments of sporadic nutrient flux and high spatial heterogeneity favor organisms that utilize resources more efficiently [3, 4]. Using batch cultures, we focus here on the latter case of pulsating environments with a single nutrient pulse where the outgoing flux of nutrients is limited so that the organism can utilize all available nutrients, including reutilization of secreted byproducts.

The biomass produced per consumed nutrient in conditions of continuous nutrient flux is physiologically defined as the biomass yield parameter that describes the efficiency of nutrient utilization [5–7]. Theoretical models, known as ‘black box models’, predict the biomass yield for growth on a single nutrient source in conditions of continuous flux such as those observed in chemostat experiments to high accuracy [8–10]. These models consider growth as a single chemical reaction with the nutrients as substrates and the produced biomass and secreted byproducts as products. By calculating the change in free energy of the whole reaction, the biomass yield is predicted. In this work, we use these models to qualitatively predict the overall biomass yield, i.e., the biomass produced per nutrient amount for conditions of limited outgoing nutrient flux where the secreted byproducts are also utilized, a condition akin to a single nutrient pulse in natural pulsating environments. We demonstrate a good fit to experimental data for batch growth of *Escherichia coli* on a single nutrient source. Since natural environments typically contain multiple nutrients[11–13], we investigate whether the overall biomass yield of a nutrient depends on the availability and metabolic properties of a second nutrient, for example if it can be degraded or only used as a building block for biomass.

The black box model describes a case without mutual effects between nutrients; hence, the overall biomass yield of each nutrient is independent of the availability of another. We tested this prediction experimentally by titrating a second nutrient to batch cultures grown on a single carbon source, demonstrating that the overall biomass yield depends not only on the availability but also on the initial amount of other nutrients, and that this mutual effect can be negative. To explain these observations, we expanded the black box model to consider whether a second nutrient can only be used for biomass synthesis or also degraded for energy generation and included mutual effects between the metabolic processes of the nutrient sources. The model qualitatively captures the experimental observations and explains how the combination of nutrients affects metabolism. Furthermore, using the model, we determine the mutual effect of different nutrient combinations on growth processes.

## Results

### Growth on a single nutrient source

The classical system to investigate the efficiency of nutrient utilization in pulsating environments where organisms have sufficient time to fully utilize all available nutrients in a single nutrient pulse are batch cultures. Here we follow growth of *E. coli* until depletion of the initial nutrient source and potential secreted byproducts when stationary phase is reached in M9 minimal medium with glucose, malate, or aspartate as sole carbon sources [14, 15]. These carbon sources were chosen as respiro-fermentative, strictly respiratory, and a degradable biomass component. The produced biomass (Δ*B*), that is the biomass reached at stationary phase minus the biomass at inoculation, was recorded as the optical density at 600nm, converted to cellular dry weight using a predetermined conversion factor [16], and plotted against the initial nutrient amount (Fig. 1; Supplementary Fig. 1). The produced biomass shows a good linear fit to the initial amount of the sole carbon source (Fig. 1B) and as such, can be describe by [14]:

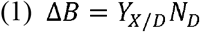

where *N_D_* is the initial amount of nutrient *D* and *Y_X/D_* the overall biomass yield on nutrient *D* which describes the efficiency of full utilization of the available nutrient.

**Fig. 1.**
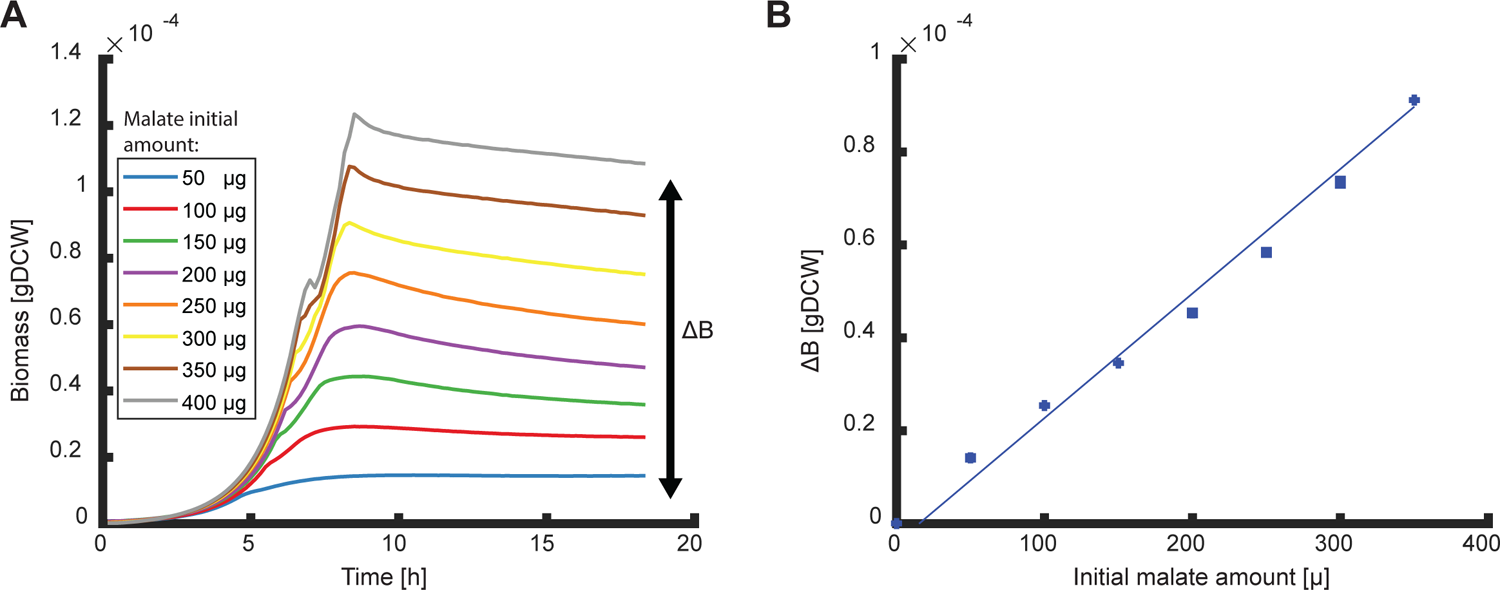
Overall biomass yield of malate. **A.** Growth curves of *E. coli* for different initial amounts of malate. Curves are averages of three biological replicates. The produced biomass (*ΔBM*) is the final biomass reached in stationary phase minus the initial biomass at inoculation. **B.** The produced biomass of the different growth curves in Fig. 1A as function of the initial nutrient amount. The slope of the linear fit is the overall biomass yield (fit parameter R^2^>0.9). Bars of standard errors of the biological replicates are too small to be noticeable. Data for glucose and aspartate experiments are shown in supplementary fig. 1.

To predict the produced biomass, we used a black box formalism [8] that describes growth of chemotrophic organisms as a two-reaction process (Fig. 2A). The first is a catabolic reaction that releases Gibbs free energy by breakdown of nutrients. The second is the anabolic reaction that uses the released free energy for the synthesis of new biomass. The overall Gibbs energy dissipation *ΔG_X_* of the growth process is given by ([8], Supplementary note A1):

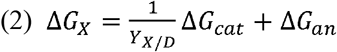

where the subscripts *cat*, and *an* refer to the Gibbs energy of dissipation of the catabolic and anabolic reactions, respectively. Given that in the here investigated growth conditions all secreted byproducts are utilized, and the free energy of the secreted byproducts can be set to 0, the overall biomass yield may be predicted as [8]:

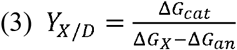

**Fig. 2.**
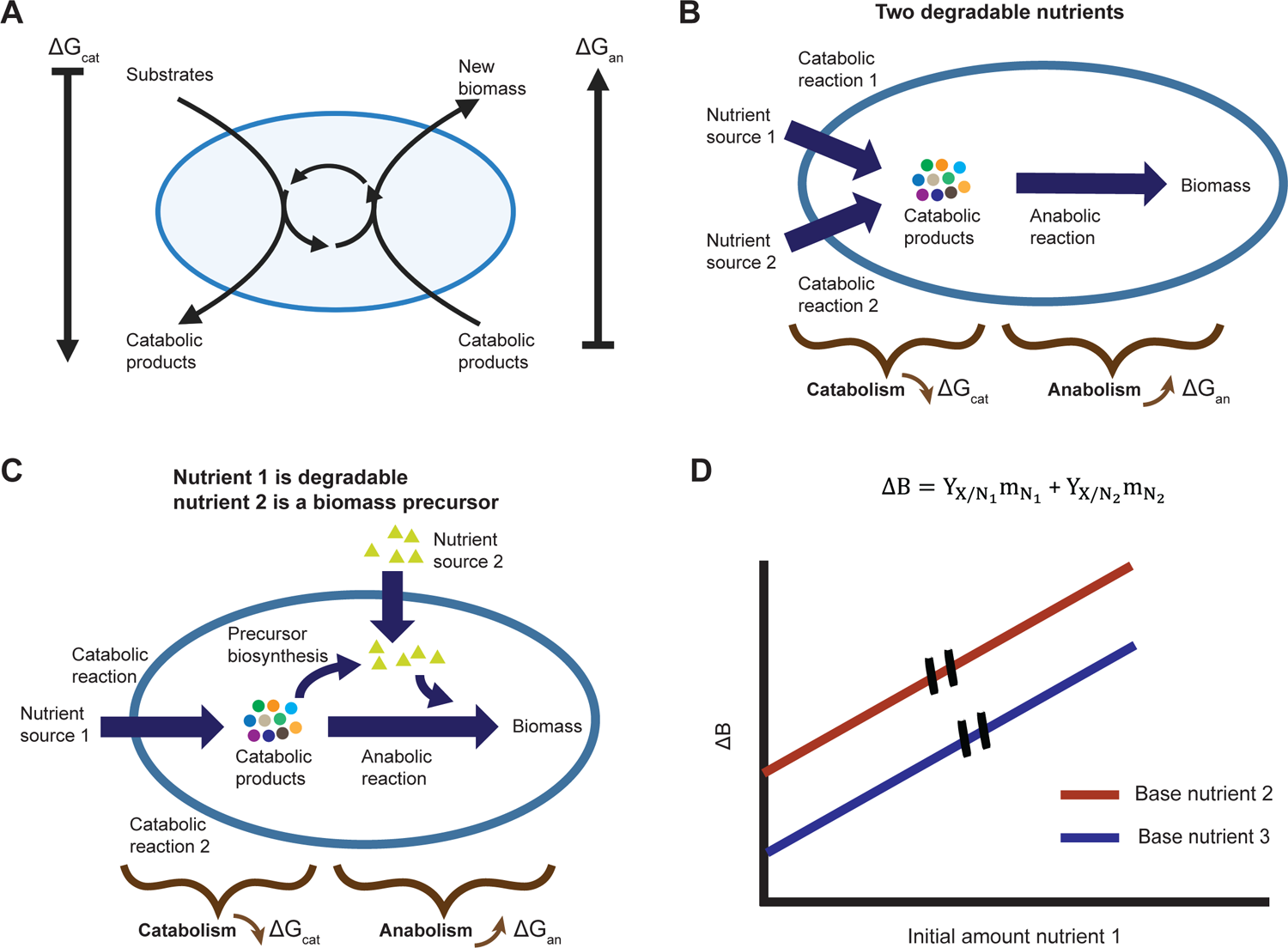
Expansion of black box model to include multiple nutrients. **A.** The overall growth process is split into two reactions – a catabolic process in which free energy is released and an anabolic process in which new biomass is formed. The ring in the middle of the cell represents the coupling of anabolism and catabolism by ATP and other biochemical process. **B.** Schematics of black box model expansion for two degradable nutrients. A catabolic reaction is added for each nutrient. **C.** Schematics of model expansion for a combination of a degradable nutrient and a second nutrient that can only be used as a biomass precursor. The anabolic reaction is separated into two reactions – one for the biosynthesis of the biomass precursor and a second for the rest of the anabolic process excluding the biosynthesis reaction of the biomass precursor. **D.** Black box model prediction for growth on two nutrient sources without mutual effect. The model predicts the produced biomass is a linear sum of the biomass gained from each nutrient. The overall biomass yield, the slope of the curve, is independent of availability of different nutrients.

Combining equations (1) and (3) predicts a linear correlation of the produced biomass as function of initial nutrient amount with a slope that depends only on the type of nutrient through *ΔG_cat_*. This prediction fits qualitatively well with all measured nutrients [14, 15] (Fig 1B, Supplementary Fig. 1).

### Growth on multiple nutrient sources

Since organisms typically encounter multiple nutrients in natural environments, we next asked whether the availability of one nutrient affects the overall biomass yield of another. To enable a black box model to capture such effects, we added another reaction that depends on the type of second nutrient: A) degradable nutrients that first must be catabolized before they can be used, such as a sugar. B) non-degradable nutrients that can be used only as a biomass precursor, such as in *E. coli* the non-degradable amino acid methionine. C) nutrients that can be both catabolized or used directly as a biomass precursor, such as in *E. coli* the amino acid aspartate. When the nutrient combination is of two degradable nutrients, the added reaction is catabolic (Fig. 2B). In this case, the overall Gibbs energy dissipation gives:

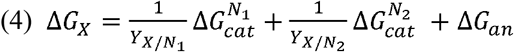

where 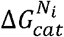 is the Gibbs energy of dissipation for the catabolic process of nutrient *i*. When the second nutrient source is a non-degradable biomass precursor, we split the anabolic reaction into two – a reaction for biosynthesis of the biomass precursor and a reaction for the general anabolic process (Supplementary A3, Fig. 2C). The overall Gibbs energy of dissipation in this case gives:

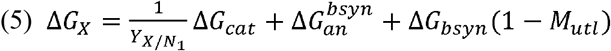

where *ΔG_bsyn_* is the Gibbs energy of dissipation for synthesis of the biomass precursor and 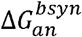 is the dissipation energy for the general anabolic process minus that of the biomass precursor. The function *M_utl_* describes the ratio of available biomass precursor to that required to generate the produced biomass during the growth process. It is dependent on biomass precursor availability such that when all the necessary biomass precursor is available in the environment, the function assumes the maximal value of 1 and the cost for this precursor biosynthesis is alleviated.

Combining equations (4) or (5) with equation (1) shows that regardless of the type of nutrient supplemented, the produced biomass is predicted to be a linear sum of the biomass gained from the available nutrients and the overall biomass yield of each nutrient is independent of the availability of others (Supplementary A2-3, Fig. 2D):

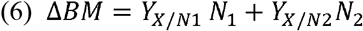

where *N_i_* is the amount of nutrient *i* in the growth medium and *Y_X/Ni_* is the overall biomass yield of nutrient *i*.

To test the prediction that the overall biomass yield of a nutrient is independent of the availability of others, we compared the overall biomass yield of *E. coli* for different nutrients, henceforth referred to as the measured nutrient, in the presence or absence of a second nutrient, termed the base nutrient. To do so, the initial amount of the measured nutrient was varied for each batch culture experiment at constant initial amounts of the base nutrient between 0 and 1.2 g/l for glucose, acetate, or aspartate and 0 and 0.06 g/l for methionine. The produced biomass was plotted against the initial amount of the measured nutrient and the overall biomass yield was determined as the slope of a linear fit of that curve (Fig. 3A,B). In combination with glucose, succinate, or acetate as base nutrients, we determined the overall biomass yield of xylose and methionine as measured nutrients, as examples of degradable or non-degradable nutrients, respectively (Fig. 3). The initial amount of base nutrient determines the intercept with the Y-axis and was chosen such that the measured parameters remain within measurable range.

**Fig. 3.**
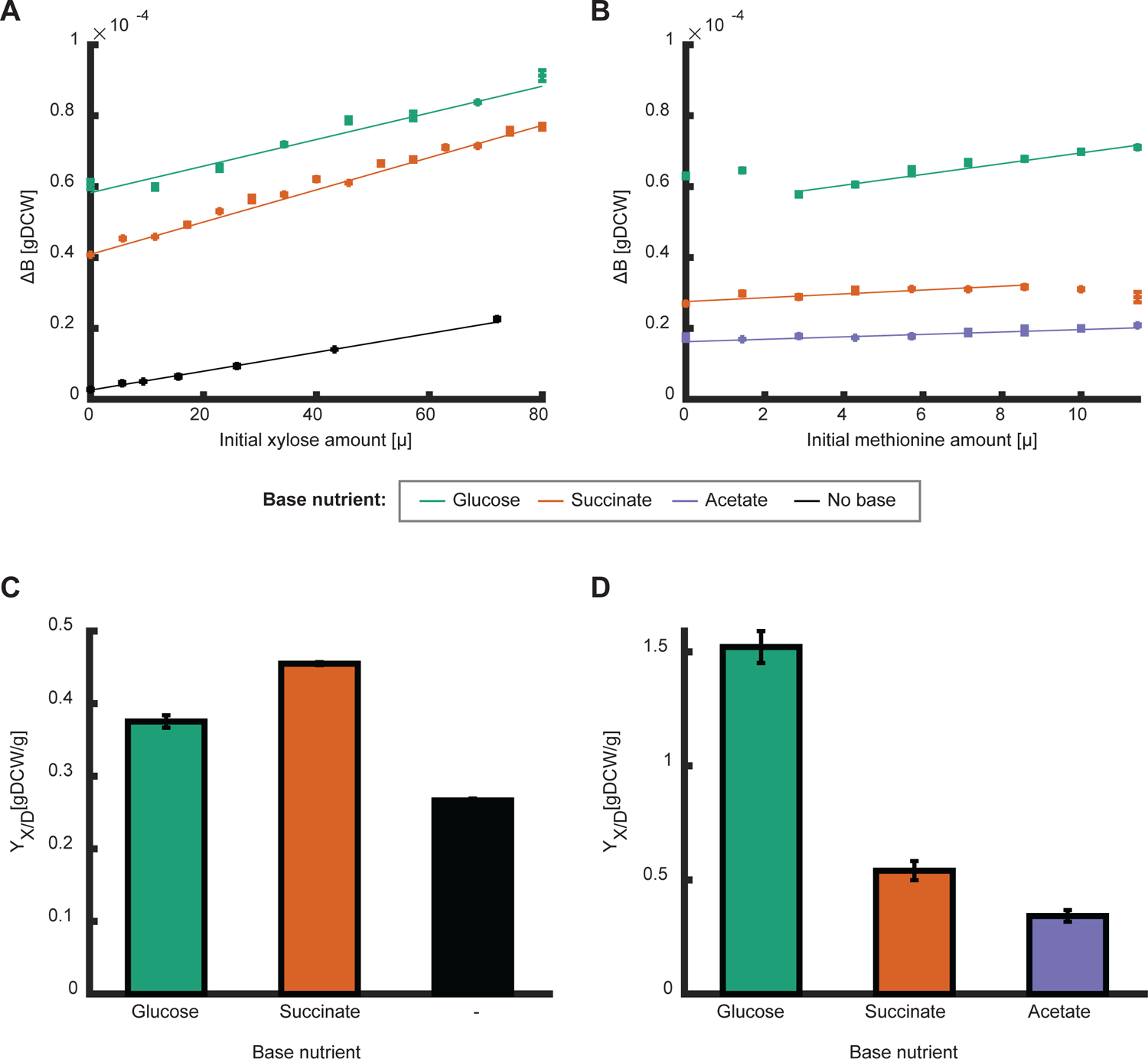
produced biomass and biomass yield on different nutrient bases. **A-B.** The produced biomass as function of initial amounts of the measured nutrients: xylose with or without different base nutrient sources (A, black– no base, green – 160 μg glucose, orange −160 μg succinate), and methionine with different base nutrient sources (B, green – 160 μg glucose, orange – 80 μg succinate, purple – 160 μg acetate). Error bars depicting standard error for three biological replicates are in several cases not visible. Curves show a linear fit to the data of the linear region (fit parameter R^2^>0.9). **C-D.** The overall biomass yield (slope of the fits above) for xylose and methionine for growth on the different base nutrients. Error bars depict error of fit parameter. The overall biomass yield is dependent on the nutrient base.

The overall biomass yield was highly dependent on the base nutrient. For xylose, the overall biomass yield was higher on succinate as base nutrient than on glucose or when used alone, and for methionine the overall yield was by far the highest on glucose (Fig. 3). For most combinations, the influence of the second nutrient was monotonous across the tested concentrations, i.e., the overall biomass yield of the measured nutrient can be determined from the slope of a linear fit (Fig. 3A,B). An exception was the non-monotonous behavior of methionine as the measured nutrient in combination with glucose as a base nutrient (Fig. 3B). At low initial amounts of methionine (below 3 μg), increasing initial amounts of methionine unexpectedly decreased the produced biomass. In the higher range of initial amounts (above 3 μg), increasing methionine initial amounts increased the produced biomass linearly.

Thus, the overall biomass yield of a measured nutrient is dependent on the base nutrient, consequently black box theory cannot capture the produced biomass of multiple nutrient sources. To enable the model to describe these mutual effects, we expand it to include such effects phenomenologically. To do so, we coupled a function that is dependent on the combination of available nutrients to the Gibbs energy dissipation of each reaction in the growth processes. For simplicity, we assumed these functions are linear to the initial nutrient amount.

As such, the overall Gibbs energy dissipation of growth on two degradable nutrient sources is described as (Supplementary A4):

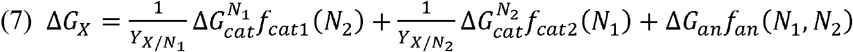

where *f_cat_*(*N_i_*), *fan*(_N_i__) are linear functions to the initial amounts of nutrient source *i*, with coefficients 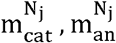 respectively. These functions phenomenologically depict the mutual effect of the nutrient combination on the growth processes. Combining equations (1) and (7) predicts the produced biomass:

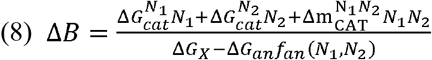

where 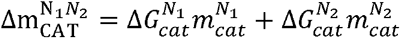 and 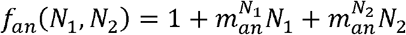. Given a mutual effect between nutrients, the produced biomass is thus made of three terms, two describing the direct effect of catabolism of the two nutrient sources and a third term describing the mutual catabolic effect depending on availability of both substrates. Depending on the type of mutualism, qualitatively different relationships are predicted between available nutrients and biomass formation (Fig. 4A) – a positive mutual catabolic effect increases the overall biomass yield (Fig 4A, orange curve) while a negative catabolic effect decreases it (Fig. 4A, purple curve). The model can capture the experimentally observed mutual effect of increased overall biomass yield with a positive mutual catabolic effect (compare increased slope for different base nutrients in Fig. 3A to the orange curve in Fig. 4A).

**Fig. 4.**
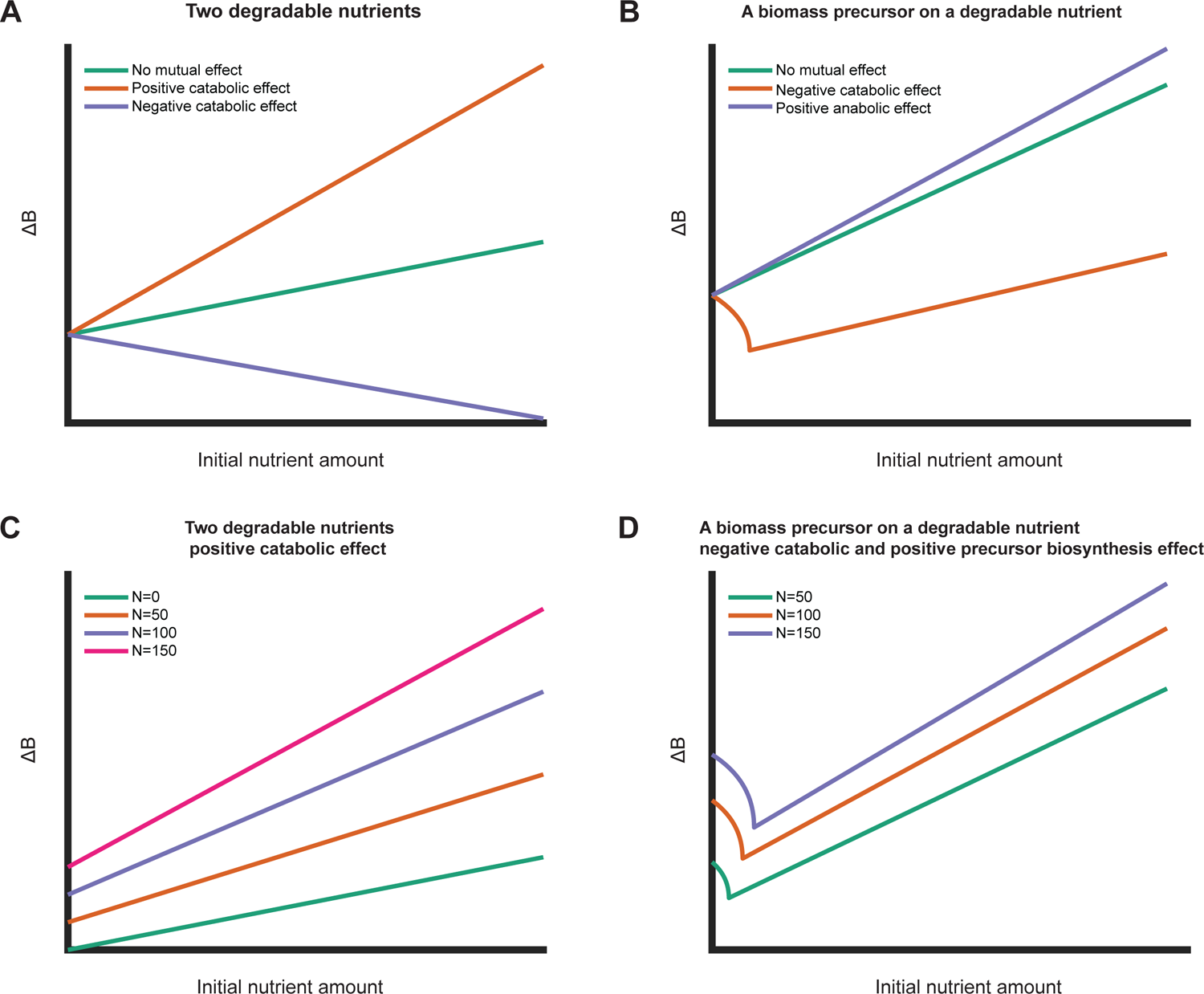
Expanded model prediction including mutual effects between two nutrients. **A-B.** Simulations of expanded model with different mutual effects for growth on two degradable nutrient sources (A) and a biomass precursor in combination with a degradable nutrient (B). Initial nutrient amount of the base nutrient were kept constant in all simulations. **C.** Simulations of expanded model for growth on two degradable nutrients for different initial amounts of base nutrient with positive catabolic mutual effect. The overall biomass yield (slope) increases with increasing initial amount of base nutrient. **D.** Simulations of expanded model for growth on a biomass precursor and a degradable base nutrient with a negative catabolic effect and positive effect on precursor biosynthesis. Increased initial amount of base nutrient shifts the initial decreasing part and increases the slope (the overall biomass yield) of the linear part.

For growth on a combination of a degradable nutrient and a non-degradable biomass precursor, the overall Gibbs energy dissipation is described as (supplementary A5):

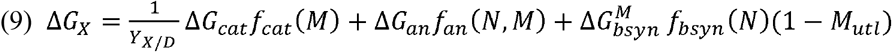

where *f_cat_*(*M_utl_*), *fan*(*N, M_utl_*), *f_bsyn_*(*M_utl_*) are linear functions with coefficients 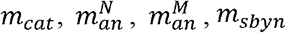 respectively. These functions depict the mutual effect between the nutrient sources on the Gibbs free energy of each growth reaction. Solving equations (1) and (9) for the produced biomass gives a quadratic equation:

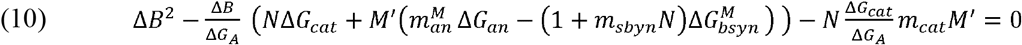

where 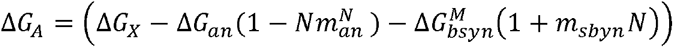 and *M*′ = *M_utl_*Δ*B*.

Solving equation (10) for the produced biomass shows that, unlike the solution for the case of growth on two degradable nutrient sources, a mutual effect between a biomass precursor and a degradable nutrient can give rise to non-monotonous solutions. Fig. 4B depicts examples of possible mutual effects between a precursor and a degradable nutrient. The case of a negative catabolic effect (Fig. 4B, orange curve) fits qualitatively well with the experimental observation of the biomass precursor methionine on glucose as base nutrient (Fig. 3B, green data points).

The fitting parameters of the linear functions are a key output of the model since they infer how each combination of nutrients effects the different growth reactions. Fitting the model to the measurement of methionine growing with glucose as a base nutrient gives a negative value for the catabolic parameter, revealing that methionine decreases the catabolic efficiency of glucose. Furthermore, the overall biomass yield of methionine on glucose in the linear region is higher than that on succinate or acetate (Fig. 3D), suggesting a mutual effect on another metabolic process in one of these combinations, potentially the anabolic process. For all combinations of two degradable nutrients, the overall biomass yield increased as compared to growth on sole nutrient sources (Fig. 3C), demonstrating a positive mutual effect on the catabolic process.

An unexpected model prediction is noticeable in equations (9) and (10) where the initial amounts of the two available nutrients are coupled in at least one term. Hence, the model predicts that the overall biomass yield of a measured nutrient depends not only on the availability of a base nutrient, but also on the relative initial amounts of the nutrients. For a combination of two degradable nutrient sources with a positive catabolic effect, as observed experimentally for xylose on the two base nutrients (Fig. 3A,C), the overall biomass yield is predicted to increase with increasing initial amounts of the base nutrient (Fig. 4C). For the combination of a degradable nutrient and a biomass precursor, such as methionine on glucose, with a negative catabolic effect and positive effect on precursor biosynthesis, the model predicts a shift of the curves for the non-linear part as well as an increase in the slope of the linear part with increasing initial amounts of base nutrient (Fig. 4D).

To test these predictions, we determined the produced biomass on xylose and methionine as the measured nutrients on different initial amounts of succinate and glucose as the base nutrients, respectively (Fig. 5A,B). The overall biomass yield of xylose (i.e., slope of the curve) increased linearly with the initial amount of the base nutrient succinate (Fig 5A, C). This observation fits well with the model prediction for a positive catabolic effect between two degradable nutrients (Fig. 4C). The curve of the produced biomass on methionine exhibits a more complex dependency on the initial amount of glucose as the base nutrient. Above 5 μg methionine, the slope of all curves increased linearly with the amount of the base nutrient glucose, but below 5 μg methionine there was no linear dependency and the amount of base nutrient varied the curve shape (Fig. 5B, D). This observation fits well with the theoretical prediction (Fig. 4D) that this nutrient combination not only has a positive effect on the precursor biosynthesis reaction (i.e., the linear dependency at higher methionine supplementation), but also a negative catabolic effect where at low methionine concentrations, in some cases, more methionine leads to lower biomass gain.

**Fig. 5.**
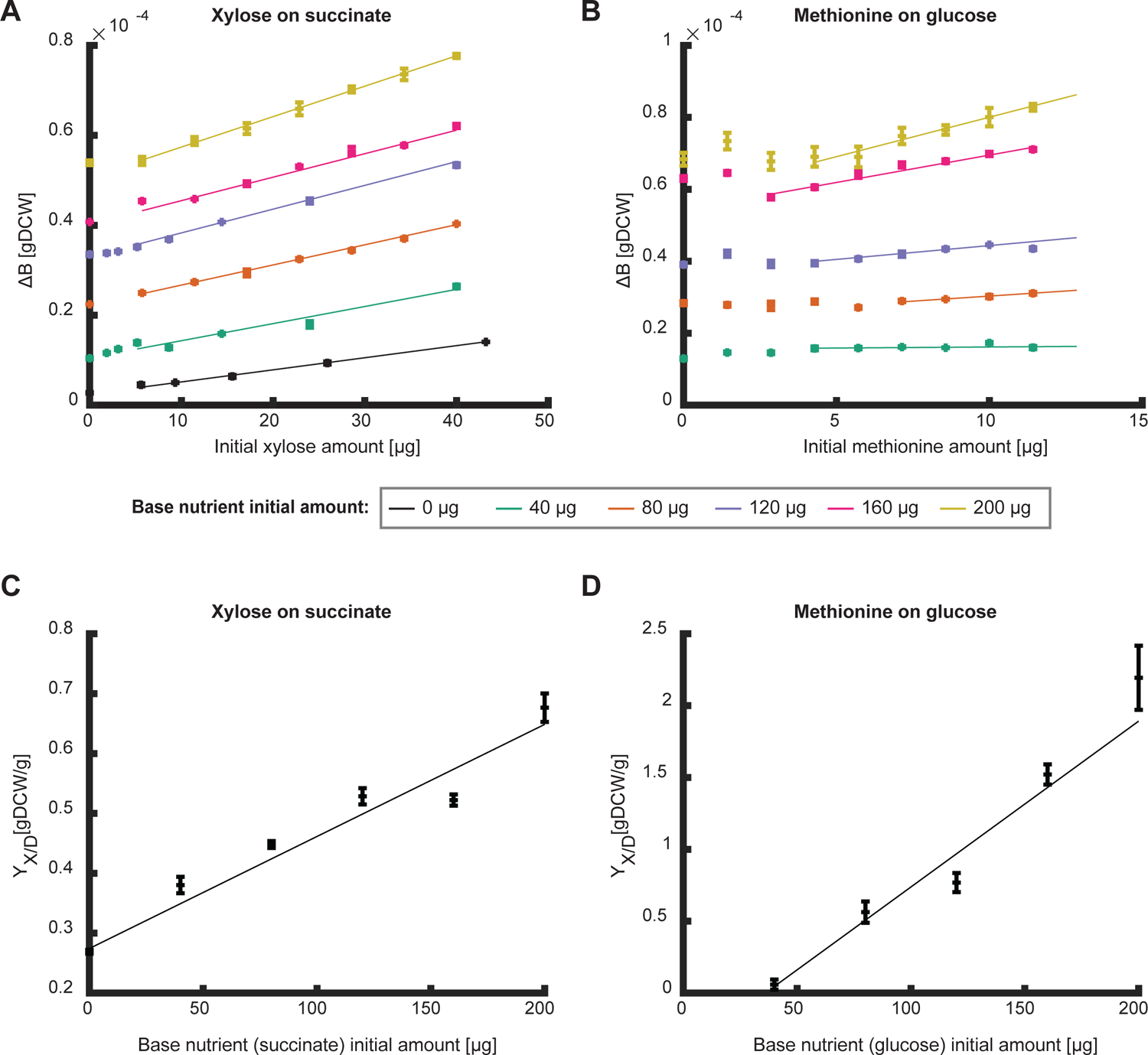
Effects of initial amounts of base nutrients on the overall biomass yield. **A-B.** The produced biomass as a function of the initial amount of xylose (A) and methionine (B) at different initial amounts of the base nutrient succinate (A) and glucose (B). Data points are average of three biological replicates and error bars are standard error that are too small to notice. Curves are linear fits in the linear region (fit parameter R^2^>0.9 for all fits except for methionine on 40 μg glucose which showed a good fit to a constant (p-value<0.05)). **C-D.** The overall biomass yield as a function of initial nutrient amount as calculate from the conditions in A-B. Curves show linear fit region (fit parameter R^2^>0.9). Error bars depict error of fit parameter.

Which mechanism underlies the negative catabolic effect of methionine on glucose? The growth curves followed the classical diauxic shift with exponential growth on glucose and a second phase on previously secreted acetate (Supplementary Fig. 2A). For the example of 160 μg glucose as the base nutrient (Fig. 5, pink curve), the first phase lasted 4-4.5 hours and growth on acetate resumed between 7-10 hours (Supplementary Fig. 2A). In both phases, the biomass gain (calculated as the biomass at the end minus the biomass at the beginning) increased linearly with methionine amounts greater than 2 μg (Supplementary Fig. 2B, C). The biomass gain was much higher than the trendline in the absence of or at very low methionine concentrations. During exponential growth on glucose in the first phase, methionine decreased the gain in biomass but increased the growth rate (Supplementary Fig. 2D). Given the diauxic shift from growth on glucose to previously secreted acetate (Supplementary Fig. 2E), the most plausible explanation for the higher biomass gain without or low methionine in the second phase is due to higher acetate secretion in the first phase. To test whether methionine supplementation indeed reduced acetate secretion, we varied acetate secretion rates by altering steady state growth through an inducible promoter for the glucose uptake gene *ptsG* that limits glucose uptake [17]. Comparing acetate secretion in the presence and absence of methionine shows that methionine indeed decreases acetate secretion (Supplementary Fig. 2F). Thus, the negative catabolic effect of methionine on glucose catabolism appears to be a combination of a lower biomass gain during the first growth phase, with a higher growth rate and less acetate secretion, and a lower biomass gain in the second phase because less acetate was secreted.

At the lowest methionine amounts (0 and 1.43 μg) we noted a shorter lag time for growth on acetate (Supplementary Fig. 2A, compare red and black curves to the other curves). Growth with 1.43 μg methionine was somewhat special as it followed the biomass trendline in the first growth phase but could not sustain the higher growth rate throughout this growth phase (Supplementary Fig. 2A, red curve between 2-4h), presumably because methionine was used up, which explains why its biomass gain in the second phase was indistinguishable from the no methionine condition (Supplementary Fig. 2C). Consistently, 1.43 μg methionine was below the amount necessary to produce the biomass reached at the end of the first growth phase (about 1.7 μg of methionine is required to generate 0.8 gDCW of biomass [19]).

So far, we focused on degradable nutrients or nutrients that can only be used as biomass precursors. Some nutrients such as degradable amino acids, however, can be directly used both as biomass precursors or energy source. Given the complex curves observed for the combination of biomass precursor and degradable nutrient, we expected that a degradable amino acid in combination with a degradable nutrient would also produce non-monotonous curves. To investigate the effects of such nutrient combinations, we measured the produced biomass on the degradable amino acid aspartate on different initial amounts of glucose and acetate as base nutrients (Fig. 6A,B). The combination of aspartate and acetate led to a complex curve with two linear phases separated by a double shift in slope at intermediate concentrations (between 100 – 150 μg, Fig. 6A). The first phase at low initial amounts of aspartate resulted in a linear slope that increases with initial amount of acetate while the slope of the second phase shows only a low dependency on acetate initial amounts (Fig. 6C). Aspartate on glucose also shows a complex curve with two linear phases (Fig. 6B). In this nutrient combination, the slope of the first phase is independent of the initial amount of glucose yet the length of this phase increases with increasing initial glucose amounts (Fig. 6D). The slope of the second linear phase increases with increasing glucose initial amounts. The complex behavior observed in these experiments cannot be captured even with the mutual effect model presented here. We hypothesize that the ratio of how much aspartate is utilized as a biomass precursor to how much is catabolized affects the overall biomass yield. The multiple utilization possibilities add an additional degree of freedom to the system and as such, capturing the behavior of these nutrients in a model requires time-resolved intracellular flux information.

**Fig. 6.**
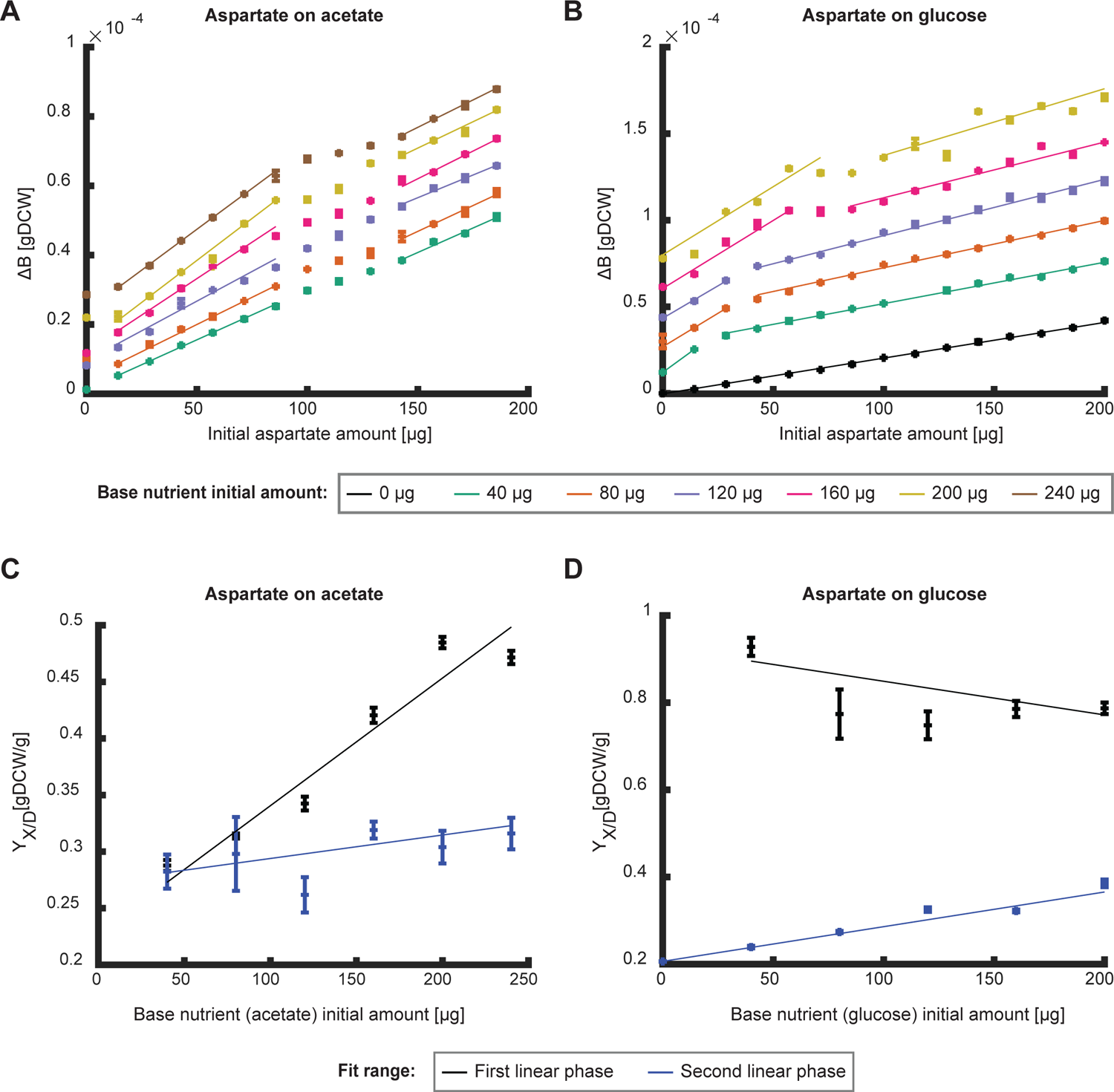
Effects of initial amounts of base nutrients on the overall biomass yield. **A-B.** The produced biomass as function of initial nutrient amount of aspartate for different initial amounts of the base nutrient (**A** – glucose, **B** – acetate). Data points are average of three biological replicates and error bars are standard error that are too small to notice. Curves are linear fits in the linear regions (fit parameter R^2^>0.9). **C-D.** The overall biomass yield as function of initial nutrient amount as calculated from the conditions in A-B respectively. Black data points show the slope of the first linear phase and blue data points show the second. Curves show linear fit (fit parameter R^2^>0.9 for first linear fit on acetate and second linear phase on glucose, the other fits show a good fit to a constant (p-value<0.05)). Error bars depict error of fit parameter.

## Discussion

In conditions of low nutrient flux, organisms utilize all nutrients in the environment and reabsorb previously secreted byproducts to fuel further growth. Here we asked whether the availability of one nutrient affects the utilization efficiency of another? We showed that different nutrient combinations have different mutual effects on an organism’s ability to generate biomass, presumably by changing intracellular metabolism, secretion, and reabsorption of secreted byproducts. To better understand these observations, we expanded previous black box models to account for the mechanistic effect of different nutrient types and phenomenologically added mutual effects between nutrient sources. The expanded black box model was able to qualitatively capture the experimental observation and further predicts that the overall biomass yield of a nutrient depends not only on the availability of other nutrients but also on the ratio of initial amounts of the different nutrients.

While the expanded model developed in this study accounts for mechanistic properties of how nutrients are utilized, its coarse granularity does not include specific metabolic reactions. Thus, the underlying molecular mechanisms remain unknown but the model demonstrates how combinations of nutrients affect the abstract metabolic reactions of catabolism, anabolism, and biomass precursor biosynthesis. By fitting the model to experimental measurements, further hypotheses can be generated about the mechanistic nature of the mutual effects between nutrients. For instance, we observed that for *E. coli* growing on glucose, methionine supplementation decreases the catabolic efficiency of glucose utilization, and provide circumstantial evidence that this is caused by a combination of the effect of methionine on the growth rate and reduced acetate secretion.

For all nutrient combinations, the initial amount of the base nutrient had a positive effect on the overall biomass yield of the measured nutrient, for at least some region of the measured range. This result is consistent with previous reports, for example, the supplementation of growth media with casamino acids or yeast extract has been shown to increase the carbon utilization efficiency of succinate or asparagine in batch culture experiments of *Enterobacter aerogenes* and *Pseudomonas perfectomarinus* [20]. Similarly, the utilization of mixtures of different dissolved organic carbon sources by bacterial communities in aquatic systems has been found to be more efficient than the utilization of a single source [21–25]. Moreover, the carbon utilization efficiency of *Candida utilis, P. oxalaticus, Saccharomyces cerevisiae, Paracoccus denitrificans*, and *Thiobacilius versutus* has been found to be higher than theoretically predicted when a nutrient source that can be utilized solely as an energy source was supplemented during balanced growth conditions [26–28]. It is tempting to conclude that the underlying molecular mechanisms for this deviation from the theoretical prediction are similar in all cases, regardless of the experimental setup and measured parameter, i.e., biomass yield as measured in a chemostat continuous culture [5–7] or the overall biomass yield as measured in batch cultures. Our analysis suggests that the molecular mechanism of this phenomenon is due to a mutual effect of the different nutrients on catabolism, although the specific metabolic pathways that are affected remain unclear. To further investigate these mechanisms and identify a possible global mechanism, a more detailed resolution of metabolic pathways and their fluxes will be necessary, possibly by combining the thermodynamic black-box approach presented here with genome-scale metabolic models [29, 30].

Microbial nutrient utilization efficiency is of great importance in various contexts, including evolution, microbiome-host interactions, synthetic biology, and climate change. Microbes metabolize a wide range of compounds, which affects the dynamics of organic matter and carbon dioxide emissions [31–33], potentially impacting agricultural productivity, ocean nutrient balance, and the global climate [33]. For example, heterotrophic microbes respire 60 gigatonnes of terrestrial organic matter annually, roughly six times the annual anthropogenic emissions. Given that in most growth environments microbes utilize multiple nutrients, understanding the mutual effects between different nutrients, as discussed in this paper, will help direct research of ecological systems and health related host-microbiome interactions. The analysis presented here was done for a single organism and experimentally tested on *E. coli*, but since the abstracted reactions occur in any metabolic system our approach can be extended to analyze growth of consortia or even larger ecological systems. It remains an open question whether there are general principles governing the here described mutual effects or whether each nutrient combination has its own unique mechanism in a given organism.

## Materials and methods

### Strains and growth essays

In the growth essays the NCM3722 strain [34, 35] as used and in the acetate secretion essay NQ1243 [17]. Each experiment was carried out in three steps: seed culture, pre-culture and experimental culture. For seed culture, one colony from fresh LB agar plate was inoculated into test tube with M9 minimal medium with 4 gr/l glucose and cultured in 37°C shaking at 350 rpm for 8-9 hours. The cell culture was then diluted to OD_600_=0.1-0.2 in pre-warmed shake flask with m9 minimal medium with the same base nutrient as the experiment and left to grow for two hours in 37°C shaking at 350 rpm (pre-culture). The cell culture was then diluted to OD_600_=0.03-0.08 in pre-warmed 96 deep well plate with 1 ml. Each well contained medium with the experimental growth conditions (M9 minimal medium with nutrients according to experiment, each condition was set in triplicates) and mixed thoroughly. 200 μl cell culture from every well was then transferred to 96 deep well transparent essay plate and placed in Tecan microplate reader (Tecan infinite M200) for growth measurement. Microplate reader was programmed to maintain temperature at 37°C, maximal shaking and measure OD_600_ every 10 minutes.

### Data analysis

The OD measured by the microplate reader was linearized using a premeasured calibration curve. Growth curves obtained in the microplate reader were compared to growth curves obtained in shake flask and were equivalent. The optical density was then converted to dry weight according to known calibration [36]. The final biomass point was recorded at 3-5 hours after maximal OD was reached. All linear fits were done by method of list-mean-square.

### Acetate secretion rate experiment

The experiment was done at 37°C shaker shaking at 350 rpm in three steps: seed culture, pre-culture and experimental culture. For seed culture, one colony from fresh LB agar plate was inoculated into test tube with M9 minimal medium with 4 gr/l glucose and cultured in 37°C shaking at 350 rpm for 8-9 hours. The culture was then diluted in pre-warmed 96 deep well plate to an OD_600_ of 0.05-0.4 so that all cultures reached exponential phase at the same time. Each growth condition in the deep well plate was run in triplicates. All conditions contained m9 minimal medium, 4 gr/l glucose and different concentrations of the inducer for the glucose uptake promoter 3methyl-benzyl. Half of the growth conditions contained 0.1 gr/l methionine. Every 30 min, 40 μl culture from every well were collected and used to measure OD_600_ using Tecan microplate reader (Tecan Infinite M200). Another 100 μl culture from every well was collected, centrifuged at 15,000 rpm, the supernatant was collected and immediately frozen.

Supernatant were used to measure acetate concentrations using Acetate assay kit (Megazyme Acetic Acid Assay Kit). The slope of the plot of acetate concentrations versus OD_600_ for all replicates (multiplied with the measured growth rate) was used to determine the acetate secretion rate.

## Supporting information

Supplementary figure description

## Acknowledgments

OGo and US acknowledge support by Marie Skłodowska-Curie Actions ITN “SynCrop” (grant agreement no. 764591). Figures were generated using Biorender.

## Author contributions

OGo conceived and designed the study, designed experiments, performed experiments, analyzed data and wrote the manuscript. OGa performed experiments. LE designed experiments. US wrote the manuscript.

## Additional information

**Supplementary information** The online version contains supplementary material

**Correspondence** and requests for materials should be addressed to Uwe Sauer.

