## Supplementary note 1 for "Overall biomass yield on multiple nutrient sources"

**Supplementary note: black box model of biomass yield with expansion for multiple nutrient sources**

In this note, we provide a full derivation of the model described in the main text. Then, we describe the parameters used in the model simulations.

A. **Model description**

We constructed a model to describe the produced biomass gained from growth on multiple nutrients. The model is based on the description of the ‘black box’ model described by Liu et al [1] in which the growth process is separated into a catabolic and anabolic reactions. We expanded this model to take into account growth on multiple nutrient sources. We investigated the effect of two types of nutrients: degradable nutrients that have to be catabolized before they can be used such as sugars, and non-degradable nutrients that can be used only as biomass precursors such as such as in *E. coli* the non-degradable amino acid methionine. Nutrients that can be used for both, directly as biomass precursors or that can be catabolized for energy first, such as degradable amino acids are not discussed in this analysis.

**A1. Growth on a single nutrient source:** Heterotrophic growth on a single nutrient source can be described as a two reaction process: catabolic and anabolic. A general form for the stoichiometry of these reactions can be written as follows:

(S1a)	Catabolism: $D+Y_{A}^{cat}A\to Y_{P}^{cat}P+Y_{DOX}^{cat}DOX \left( \Delta G_{cat} \right)$

(S1b)	Anabolism: $Y_{P}^{an}P+Y_{DOX}^{an}DOX+Y_{NS}^{an}NS\to X+Y_{A}^{an}A \left( \Delta G_{an} \right)$

where $D, A, NS, X, P$ and $DOX$ are electron donor (energy source), electron acceptor, nitrogen donor, dry biomass, reduced electron acceptor (product) and the oxidized electron donor, respectively. Dividing equation (S1a) by $Y_{X/D}$ and adding the Gibbs free energy of the reaction to that of reaction (S1b) gives the overall Gibbs energy change of the growth reaction [1]

(S2)	$\Delta G_{X}=\frac{1}{Y_{X/D}}\Delta G_{cat}+\Delta G_{an}$

where $\Delta G_{X}$ denotes the overall standard Gibbs energy change or standard Gibbs energy dissipation generated by the growth reaction. Assuming that no byproducts are secreted, $\Delta G_{cat}$ and $\Delta G_{an}$ can be computed from thermodynamic tables of the combustion energy of the electron donor and that of dry biomass. As such, the biomass yield for growth on a single nutrient can be predicted by:

(S3)	$Y_{X/D}=\frac{\Delta G_{cat}}{\Delta G_{X}-\Delta G_{an}}$

**A2. Growth on two degradable nutrient sources without mutual effect:** Here we look at the model prediction for a combination of a two degradable nutrients that have to be catabolized to utilize. To account for a second nutrient source that has to be catabolized, we add a second catabolic reaction (fig. 2B of main text). The full growth process can then be written as follows:

(S4a)	Catabolism 1: $D_{1}+Y_{A}^{cat1}A\to Y_{P}^{cat1}P+Y_{DOX}^{cat1}DOX \left( \Delta G_{cat}^{N_{1}} \right)$

(S4b)	Catabolism 2: $D_{2}+Y_{A}^{cat2}A\to Y_{P}^{cat2}P+Y_{DOX}^{cat2}DOX \left( \Delta G_{cat}^{N_{2}} \right)$

(S4c)	Anabolism: $Y_{P}^{an}P+Y_{DOX}^{an}DOX+Y_{NS}^{an}NS\to X+Y_{A}^{an}A \left( \Delta G_{an} \right)$

equations (S4a) and (S4b) describe the catabolic process of nutrient $N_{1}$ (electron donor $D_{1}$) and $N_{2}$ (electron donor $D_{2}$), respectively. Dividing the two catabolic equations by the respective yield of each process and summing equation (S5a – S4c) gives:

(S5)	$\Delta G_{X}=\frac{1}{Y_{X/N_{1}}}\Delta G_{cat}^{N_{1}}+\frac{1}{Y_{X/N_{2}}}\Delta G_{cat}^{N_{2}}+\Delta G_{an}$

The overall biomass yield of nutrient $i$ is defined as (equation (1) in main text):

(S6)	$Y_{X/N_{i}}=\frac{\Delta B}{N_{i}}$

where $\Delta BM$ is the produced biomass and $N_{i}$ is the initial amount of nutrient $i$. Combining equations (S5) and (S6) and solving for the produced biomass gives:

(S7)	$\Delta B=\frac{\Delta G_{cat}^{N_{1}}}{\left( \Delta G_{X}-\Delta G_{an} \right)}N_{1}+\frac{\Delta G_{cat}^{N_{2}}}{\left( \Delta G_{X}-\Delta G_{an} \right)} N_{2}$The produced biomass is predicted to be linear to the initial amount of each nutrient with a slope that is independent of availability of the other nutrient.

**A3. Growth on a degradable nutrient and a second non-degradable nutrient that can be used only as a biomass precursor:** Here we look on the model prediction for a combination of a degradable nutrient source that has to be catabolized and a nutrient source that can be used only as a biomass precursor ($M$) such as a non-degradable amino acid. To account for this combination we separate the anabolic reaction into two reactions. One reaction describes the biosynthesis of the available metabolite and the second reaction describes the overall anabolic reaction excluding the reaction for the biosynthesis of the available metabolite (fig. 2A of main text):

(S8a)	Catabolism: $D+Y_{A}^{cat}A\to Y_{P}^{cat}P+Y_{DOX}^{cat}DOX \left( \Delta G_{cat} \right)$

(S8b)	Metabolite biosynthesis: $Y_{p}^{bsyn}P+Y_{DOX}^{bsyn}DOX+Y_{NS}^{bsyn}NS\to Y_{bsyn}M \left( \Delta G_{bsyn} \right)$

(S8c)	Anabolism: $Y_{P}^{an}P+Y_{DOX}^{an}DOX+Y_{NS}^{an}NS+Y_{M}^{an}M\to X+Y_{A}^{an}A \left( \Delta G_{an}^{bsyn} \right)$

In the case in which the biomass precursor is not available in the environment, the cell must biosynthesize all of the precursor and the overall Gibbs energy change of the reaction gives:

(S9) $\Delta G_{X}=\frac{1}{Y_{X/D}}\Delta G_{cat}+\Delta G_{an}+\Delta G_{bsyn}$

Solving eq. (S9) for the biomass yield gives:

(S10) $Y_{X/D}=\frac{\Delta G_{cat}}{\Delta G_{X}-\left( \Delta G_{an}+\Delta G_{bsyn} \right)}$

This is the exact same solution as equation (S3) except that here, the anabolic process is separated into two parts, anabolism and precursor biosynthesis.

We next examine the effect of supplementing the metabolite to the growth media. When supplemented, the cell is able to uptake the metabolite and alleviate the cost for the biosynthesis of this precursor. As such, the overall Gibbs energy change of the reaction gives:

(S11) $\Delta G_{X}=\frac{1}{Y_{X/D}}\Delta G_{cat}+\Delta G_{an}^{bsyn}+\Delta G_{bsyn}(1-M_{utl})$

where $M_{utl}$ is the portion of precursor utilized from the environment out of the total amount of precursor used in the entire growth process. Assuming all the precursor available in the environment is utilized:

(S12) $M_{utl}=\frac{M_{supp}}{M_{tot}}$

where $M_{supp}$ is the amount of supplemented precursor and $M_{tot}$ is the total amount of precursor that was used in the entire growth process coming from either precursor biosynthesis or from the environment. Given that there is a fixed amount of the precursor necessary for a growth reaction, $M_{reac}$, the total amount of precursor used in the reaction is:

(S13) $M_{tot}=M_{reac}*\Delta B$

Combining equations (S9), (S11-S13) and solving for the produced biomass gives:

(S14)$\Delta BM=\frac{\Delta G_{cat}}{\left( \Delta G_{X}-\Delta G_{an}-\Delta G_{bsyn} \right)}N+\frac{\Delta G_{bsyn}}{\left( \Delta G_{X}-\Delta G_{an}-\Delta G_{bsyn} \right)}\frac{M_{supp}}{M_{reac}}$

The produced biomass is linear to both the initial amount of $N$ and the supplemented precursor $M_{supp}$ with a slope that is independent of availability of other nutrients.

**A4. Growth on two degradable nutrient sources with mutual effect:** The theoretical prediction, given above, of a produced biomass that is a linear sum of the available nutrients doesn’t fit the experimental results which shows a mutual effect between nutrients (fig. 3 in main text). To account for this, we phenomenologicaly add an effect to the Gibbs dissipation energy of each reaction in the growth process that is based on the availability other nutrients. While the substrates and products of each reaction stay the same, the Gibbs energy change of each reaction is varies according to the availability of other nutrients such that, for the case of growth on degradable nutrients, the grow reactions are:

(S15a)	Catabolism 1: $D_{1}+Y_{A}^{cat1}A\to Y_{P}^{cat1}P+Y_{DOX}^{cat1}DOX \left( \Delta G_{cat}^{N_{1}} f_{cat1}\left( N_{2} \right) \right)$

(S15b)	Catabolism 2: $D_{2}+Y_{A}^{cat2}A\to Y_{P}^{cat2}P+Y_{DOX}^{cat2}DOX \left( \Delta G_{cat}^{N_{2}} f_{cat2}\left( N_{1} \right) \right)$

(S15c)	Anabolism: $Y_{P}^{an}P+Y_{DOX}^{an}DOX+Y_{NS}^{an}NS\to X+Y_{A}^{an}A \left( \Delta G_{an}f_{an}\left( N_{1},N_{2} \right) \right)$

where $f_{cat1}, f_{cat2}$ and $f_{an}$ are functions that depict the mutual effect of the nutrients on the reactions of equations (S4a), (S4b) and (S4c) respectively. The mutual effect on the catabolic reactions is dependent on the availability of the second nutrient while the effect on the anabolic function is dependent on availability of both nutrients. The overall Gibbs energy change of the growth reaction gives:

(S15)	$\Delta G_{X}=\frac{1}{Y_{X/N_{1}}}\Delta G_{cat}^{N_{1}}f_{cat1}\left( N_{2} \right)+\frac{1}{Y_{X/N_{2}}}\Delta G_{cat}^{N_{2}}f_{cat2}\left( N_{1} \right)+\Delta G_{an}f_{an}\left( N_{1},N_{2} \right)$

We assume that the mutual effect functions are linear to the nutrient amount such that: $f_{cat1}\left( N_{2} \right)=1+m_{cat}^{N_{2}}N_{2}$, $f_{cat2}\left( N_{1} \right)=1+m_{cat}^{N_{1}}N_{1}$ and $f_{an}\left( N_{1}, N_{2} \right)=1+m_{an}^{N_{1}}N_{1}+m_{an}^{N_{2}}N_{2}$. Solving equation (S15) for the produced biomass gives:

(S16)	$\Delta B=\frac{\Delta G_{cat}^{N_{1}}N_{1}+\Delta G_{cat}^{N_{2}}N_{2}+\left( \Delta G_{cat}^{N_{1}}m_{cat}^{N_{1}}+\Delta G_{cat}^{N_{2}}m_{cat}^{N_{2}} \right)N_{1}N_{2}}{\Delta G_{X}-\Delta G_{an}f_{an}\left( N_{1}, N_{2} \right)}$

The parameters $m_{cat}^{N_{1}}$, $m_{cat}^{N_{2}}$, $m_{an}^{N_{1}}$and $m_{an}^{N_{2}}$ describe the effect of the nutrients on the other reactions and can get any real value. Figure 4A,C in the main text shows the model prediction for the produced biomass for different parameter values.

**A5. Growth on a degradable nutrient source and a second non-degradable nutrient that can be used only as a biomass precursor with mutual effect:** Similar to the case presented in supplementary section A4, we now add a mutual effect between the two nutrient sources for the case of growth on a degradable nutrient source and a biomass precursor such as a non-degradable amino acid. We again add a phenomenological effect to the Gibbs dissipation energy that is based on the availability other nutrients for each reaction. In this case the growth reactions are:

(S17a)	Catabolism: $D+Y_{A}^{cat}A\to Y_{P}^{cat}P+Y_{DOX}^{cat}DOX \left( \Delta G_{cat}f_{cat}\left( M \right) \right)$

(S17b)	Metabolite biosynthesis: $Y_{p}^{bsyn}P+Y_{DOX}^{bsyn}DOX+Y_{NS}^{bsyn}NS\to Y_{bsyn}M \left( \Delta G_{bsyn}^{M} f_{sbyn}\left( N \right) \right)$

(S17c)	Anabolism: $Y_{P}^{an}P+Y_{DOX}^{an}DOX+Y_{NS}^{an}NS+Y_{M}^{an}M\to X+Y_{A}^{an}A \left( \Delta G_{an}f_{an}\left( N,M \right) \right)$

where $f_{cat}\left( M \right)$, $f_{sbyn}\left( N \right)$ and $f_{an}\left( N,M \right)$ are function depicting the mutual effect between nutrients and are dependent on availability of other nutrients in the growth media. The Gibbs dissipation energy of the whole growth process gives:

(S18) $\Delta G_{X}=\frac{1}{Y_{X/D}}\Delta G_{cat}f_{cat}\left( M \right)+\Delta G_{an}f_{an}\left( N,M \right)+\Delta G_{bsyn}^{M} f_{bsyn}\left( N \right)(1-M_{utl})$

Again, the energy cost for biosynthesis of the metabolic precursor is alleviated by utilization of the precursor from the environment as explained in supplementary section A3. We assume again that the mutual effect function are linear: $f_{cat}\left( M_{utl} \right)=1+m_{cat}M_{utl}$, $f_{bsyn}\left( N \right)=1+m_{sbyn}N$ and $f_{an}\left( N,M_{utl} \right)=1+m_{an}^{N}N+m_{an}^{M}M_{utl}$. Combining equations (S18), (S12), (S13) and (S9) gives a quadratic equation:

(S19) $\Delta B^{2}-\Delta B\left( N\Delta G_{C}+\frac{M_{supp}}{M_{reac}}\left( m_{an}^{M}\Delta G_{A}-\left( 1+m_{sbyn}N \right)\Delta G_{BSYN} \right) \right)-N\Delta G_{C}m_{cat}\frac{M_{supp}}{M_{reac}}=0$

Solving equation (S19) for $\Delta BM_{1,2}$gives:

(S20)

$$\Delta B_{1,2}=\frac{1}{2}\left( N\Delta G_{C}+\frac{M_{supp}}{M_{reac}}\left( m_{an}^{M}\Delta G_{A}-\left( 1+m_{sbyn}N \right)\Delta G_{BSYN} \right) \right)\pm\frac{1}{2}\sqrt{\left( N\Delta G_{C}+\frac{M_{supp}}{M_{reac}}\left( m_{an}^{M}\Delta G_{A}-\left( 1+m_{sbyn}N \right)\Delta G_{BSYN} \right) \right)^{2}+4\Delta G_{C}m_{cat}N\frac{M_{supp}}{M_{reac}}}$$

where:

$\Delta G_{C}=\frac{\Delta G_{cat}}{\left( \Delta G_{X}-\Delta G_{an}-N\Delta G_{an}m_{an}^{N}-\left( 1+m_{sbyn}N \right)\Delta G_{bsyn}^{M} \right)}$, $\Delta G_{A}=\frac{\Delta G_{an}}{\left( \Delta G_{X}-\Delta G_{an}-N\Delta G_{an}m_{an}^{N}-\left( 1+m_{sbyn}N \right)\Delta G_{bsyn}^{M} \right)}$, $\Delta G_{BSYN}=\frac{\Delta G_{bsyn}^{M}}{\left( \Delta G_{X}-\Delta G_{an}-N\Delta G_{an}m_{an}^{N}-\left( 1+m_{sbyn}N \right)\Delta G_{bsyn}^{M} \right)}$.

Equation (S20) shows that the produced biomass is predicted to be non-monotonous to initial nutrient concentrations depending on the values of the mutual effect parameters $m_{cat}$, $m_{sbyn}$, $m_{an}^{N}, m_{an}^{M}$. Figure 4B,D in the main text shows the model prediction for the produced biomass for different parameter values.

**A5. Simulation parameters**

The simulation parameters for Figure 4 are calculated according to the dissipation energy of the nutrient[1, 2]: $\Delta G=-86.6-94.4\gamma$

Where $\gamma$ is the degree of reduction of the compound using as reference state CO_2_, H_2_O and nitrogen in its most oxidized form in which it occurs in the respective growth system.

$\Delta G_{cat}^{N_{1}}=-1975 kJ C-mol^{-1}$ (xylose)

$\Delta G_{cat}^{N_{2}}=-2352 kJ C-mol^{-1}$ (glucose)

$\Delta G_{BSYN}=2920 kJ C-mol^{-1}$ (methionine)

The Gibbs energy of the overall growth reaction and the anabolic reaction [1, 3]:

$\Delta G_{X}=-3500 kJ C-mol^{-1}$

$\Delta G_{an}=500 kJ C-mol^{-1}$

The ration of available biomass precursors to that required to generate the produced biomass is calculated as [4, 5] :

$M_{reac}=ratio of methionine in proteome * part of proteins out of total biomass weight=0.0144$

Simulation parameters figure 5:

$$m_{cat}=-1$$

$$m_{sbyn}=0.01$$
