## Supplementary figure description for "Overall biomass yield on multiple nutrient sources"

**Supplementary fig. 1 – Overall biomass yield for a single nutrient**

**A-B.** Examples of growth curves of *E. coli* for different initial amounts of glucose (A) and aspartate (B). Curves are averages of three biological replicates. **C-D.** The produced biomass of the different growth curves in Supp. fig. 1A (C) and Supp. fig. 1A (D) as function of the initial nutrient amount. The slope of the linear fit is the overall biomass yield (fit parameter R^2^>0.9). Bars of standard errors of the biological replicates are too small to be noticeable.

**Supplementary fig. 2 – physiological effects in the combination of methionine and glucose**

**A.** Examples of growth curves of *E. coli* for different initial amounts of methionine on 160 $\mu g$ glucose as the base nutrient. **B.** Produced biomass during the first growth phase of the different growth curves in Supp. fig. 2A as function of the initial nutrient amount. Bars represent standard errors of three biological replicates. **C.** Produced biomass during the second growth phase of the different growth curves in Supp. fig. 2A as function of the initial nutrient amount. Values are calculated as the final biomass reached minus the biomass at the end of the first growth phase. Bars represent standard errors of three biological replicates. **D.** Growth rate during the first growth phase with methionine (between $1.43-11.43 \mu g$) and without methionine supplementation on 160 $\mu g$ glucose as the base nutrient. Methionine supplementation increased the growth rate at all amounts of glucose between 0.08 and 0.16h^-1^ (data now shown). **E.** Biomass and acetate concentration as function of time for growth on 160 $\mu g$ glucose. Error bars depict standard deviation from three technical replicates. **F.** Acetate secretion rate as function of growth rate in glucose batch cultures. The growth rate was controlled via an inducible promoter for the glucose uptake gene *ptsG*.
