## Supplementary figures and images for "Overall biomass yield on multiple nutrient sources"

### Supplementary figure 1

Supplementary figure 1

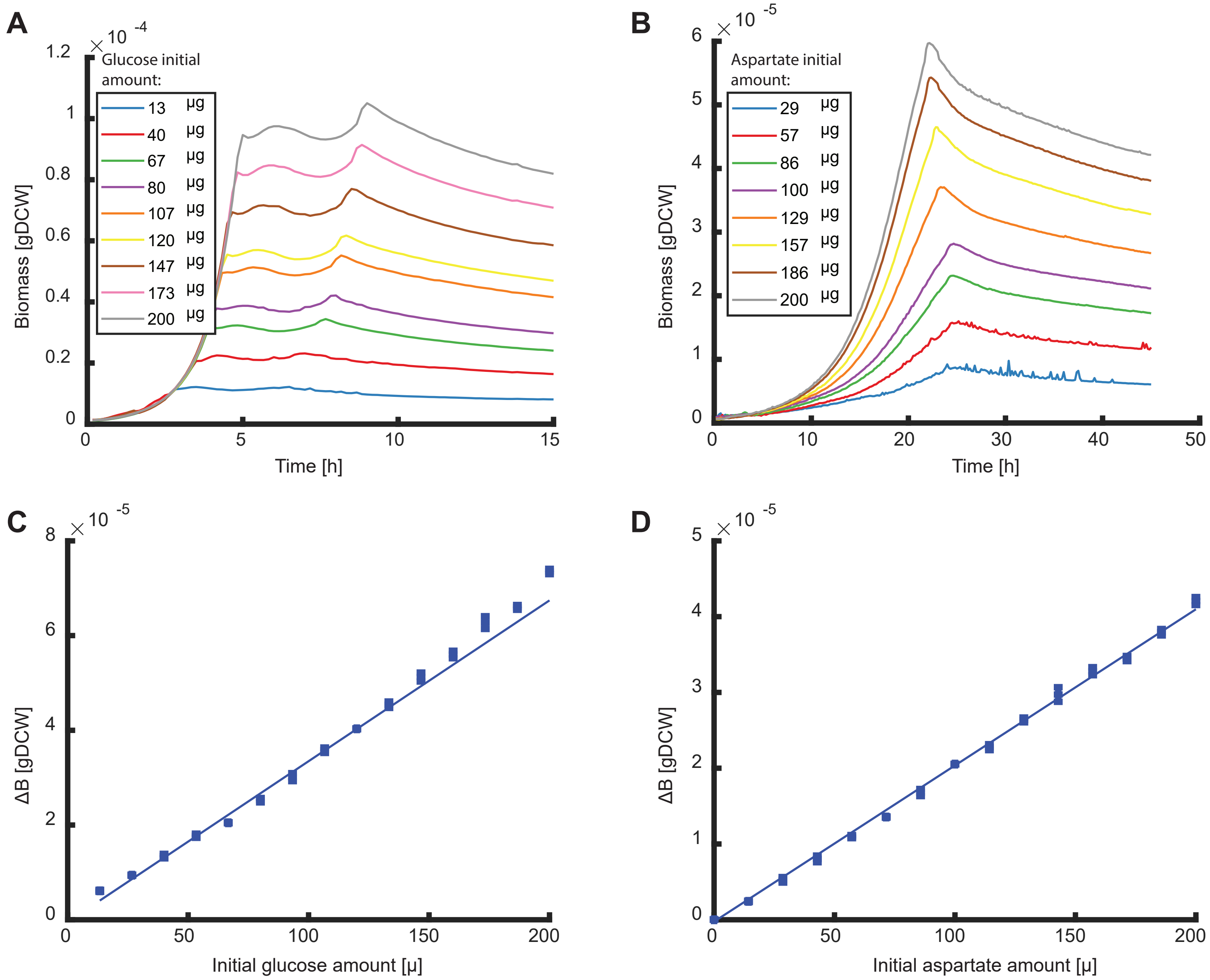

### Supplementary figure 2

#### Supplementary figure 2

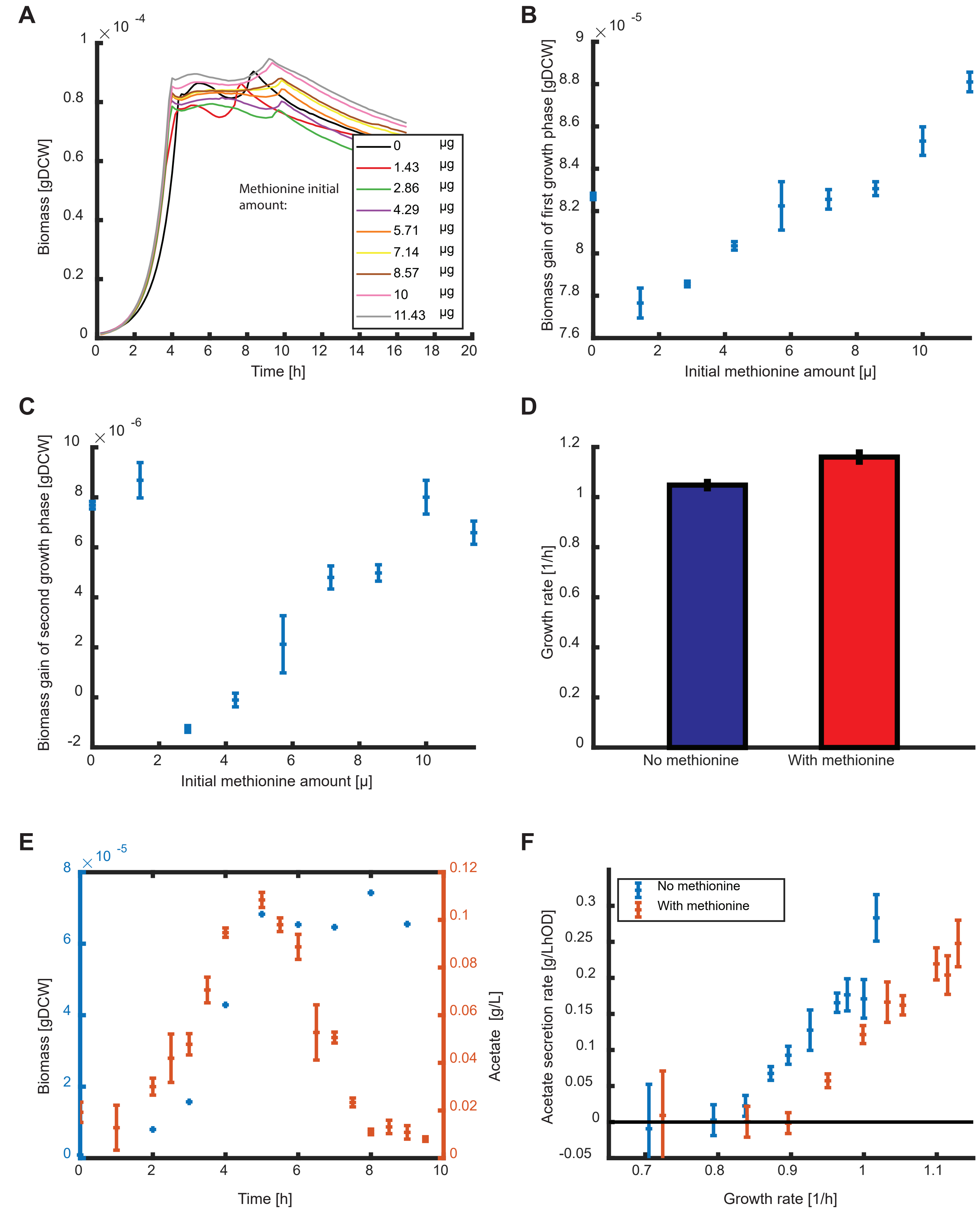
